# Non-canonical role for *Lpar1-EGFP* subplate neurons in early postnatal somatosensory cortex

**DOI:** 10.1101/2020.05.12.088450

**Authors:** Filippo Ghezzi, Andre Marques-Smith, Paul Anastasiades, Daniel Lyngholm, Cristiana Vagnoni, Alexandra Rowett, Anna Hoerder-Suabedissen, Yasushi Nakagawa, Zoltán Molnár, Simon Butt

**Author notes:** These authors contributed equally to the manuscript (Filippo Ghezzi, Andre Marques-Smith, Simon Butt). These authors contributed equally to the manuscript (Paul Anastasiades, Daniel Lyngholm, Cristiana Vagnoni).

## Abstract

Subplate neurons (SPNs) are a transient neuronal population shown to play a key role in nascent sensory processing relaying thalamic information to the developing cerebral cortex. However there is little understanding of how heterogeneity within this population relates to emergent function. To address this question we employed optical and electrophysiological technologies to investigate the synaptic connectivity of SPNs defined by expression of the *Lpar1-EGFP* transgene through the first postnatal week in primary whisker somatosensory cortex (S1BF) in mouse. Our data identify that the *Lpar1-EGFP* SPNs represent two morphological subtypes: (1) transient, fusiform SPNs with axons largely restricted to the subplate zone; (2) pyramidal SPNs with axon collaterals that traverse the overlying cortex to extend through the marginal zone. Laser scanning photostimulation of caged glutamate was used to determine columnar glutamatergic and GABAergic input onto both of these SPN subtypes. These experiments revealed that the former receive translaminar input from more superficial cortical layers up until the emergence of the whisker barrels (~postnatal (P)5). In contrast, pyramidal SPNs only receive local input from the adjacent subplate network at early ages but then at later ages can acquire varied input from the overlying cortex. Combined electrical stimulation of the ventral posterior nucleus of the thalamus and optogenetic activation of thalamic afferents in thalamocortical slice preparations revealed that *Lpar1-EGFP* SPNs only receive sparse thalamic innervation during early postnatal development. Taken together, these data reveal two components of the postnatal network that interpret sparse thalamic input to direct the emergent columnar structure of neonatal somatosensory cortex.

## INTRODUCTION

The emergence of function in the developing mammalian cerebral cortex is dependent on a diverse range of genetic and physiological processes that sculpt emergent network architecture. Fundamental research in animal models has revealed that transient neuronal circuits, observed in a restricted time window during early postnatal development, are a common feature of many cortical areas ^1, 2^. One of the first such transient circuits to be identified was that between subplate neurons (SPNs) and thalamo-recipient spiny stellate cells in layer L(4); a circuit demonstrated to play a role in the maturation of thalamocortical synapses ^1, 3, 4^. The subplate is a transient layer in the developing neocortex located between the emergent cortical plate and the underlying white matter ^5, 6^. It contains a diverse population of neuronal subtypes that differ in term of molecular markers ^7^, morphology ^8^, neurotransmitter identity^9^, and connectivity^10^. Electrophysiological studies performed in primary sensory areas suggest that SPNs are relatively mature when compared to cortical neurons in the more superficial cortical plate early in development ^8, 11^. As such they are regarded as key mediators of early spontaneous and sensory-evoked activity^4^, and direct circuit maturation^1^. A large proportion of SPNs disappear during the first postnatal week in the mouse cortex^7^. SPNs that do not undergo programmed cell death continue to form a thin, compact sublayer of layer 6, termed layer (L)6b in mature neocortex^12, 13^. Layer 6b can be cytoarchitectonically distinguished from embryonic stages.

The canonical role of SPNs is to support the establishment of thalamocortical synapses onto L4 neurons^1, 3^. In support of this model, previous studies have reported that SPNs receive thalamocortical input prior to innervation of L4 neurons ^3, 14, 15, 16, 17^. In turn, SPNs are proposed to form feed-forward connections onto thalamo-recipient L4 neurons in a transient circuit that is eliminated upon establishment of the mature thalamocortical connectivity in L4 ^10, 18^. It was proposed that, by relaying thalamic inputs to L4 through SPNs, this connectivity pattern supports developmental plasticity mechanisms ^19^ prior to the appearance of the definitive cortical architecture in primary sensory areas e.g. ocular dominance in primary visual cortex (V1) ^3^, barrel field formation in primary somatosensory cortex (S1BF) ^4^, and tonotopic organization in the primary auditory cortex (A1) ^20^. In parallel, SPNs also promote the maturation of cortical GABAergic neurons ^19^, pioneer cortico-thalamic projections ^21^, secrete proteins that control extracellular matrix composition, attract and guide thalamocortical fibres, regulate plasticity and myelination ^22^, and control the radial migration of cortical neurons at embryonic ages ^23^.

However a number of unresolved questions remain regarding SPN function in neonatal cortex: first, it remains unclear how physiological, morphological and molecular heterogeneity of subplate neurons contributes to these various roles. While previous studies performed in A1 have identified two distinct physiological populations of SPN: those that receive feedback glutamatergic input from L4 and a second cohort that only receives local input ^10^, this has not been explicitly linked to SPN identity *per se*. To this end, we have focused on a specific, genetically-defined SPN population - labelled by the *Lpar1-EGFP* transgene^7^ to understand to what extent this population represents a homogeneous subtype of SPN and better resolve the role of these cells in neonatal somatosensory cortex. Moreover, recent evidence suggests that while thalamic input is a determinant of columnar organisation in late embryonic somatosensory cortex^24^, such activity pre-dates the transition to the mature cytoarchitecture and columnar signalling unit^25^. We sought to understand the role that *Lpar1-EGFP* SPN circuits have in interpreting such information through the first week of postnatal life up until the end of the layer 4 critical period for plasticity (CPP) at around postnatal day (P)8 in the barrel field of mouse primary somatosensory cortex (S1BF). We demonstrated that *Lpar1-EGFP* SPNs represent two distinct subtypes: (1) transient (<P5) fusiform SPNs that receive columnar input from the more superficial cortical plate but whose axons and therefore output are restricted to the SP syncytium; (2) pyramidal SPNs that are found throughout the time period recorded (≤P8), that only receive local input from the SP network prior to P5 but whose axons traverse the full extent of the cortical plate to ramifying extensively through the marginal zone. Finally, we identify that thalamic input onto *Lpar1-EGFP* SPNs in S1BF is sparse throughout early postnatal life. We propose that fusiform *Lpar1-EGFP* SPNs are ideally placed to interpret and amplify sparse thalamic input alongside emergent signalling from the cortical plate, thereby providing a template - through their innervation of other SPNs including the *Lpar1-EGFP* pyramidal subtype - for the columnar circuit assembly up until the emergence of whisker barrels in L4 at ~P4-P5. Our data suggest that *Lpar1-EGFP* SPNs do not adhere to the canonical role reported for SPNs in primary sensory cortex, and supports the idea that SPNs have a variety of ways of assisting cortical circuit construction.

## RESULTS

### Intrinsic electrophysiological and morphological diversity of *Lpar1-EGFP* subplate neurons

*Lpar1-EGFP* SPNs form a layer of 2-3 cells deep adjacent to the white matter tract in neonatal S1BF (**Figure 1a**). As a first pass to understanding the contribution of these neurons to neonatal circuits of S1BF we recorded the intrinsic electrophysiological profiles of 103 SPNs from postnatal day (P)1 to P8. By injecting both depolarizing and hyperpolarizing current steps (500ms) we established that recorded SPNs had an intrinsic electrophysiological profile consistent with regular firing pyramidal cells (**Figure 1b,c**). Analysis of passive (**Figure 1d-f**) and active (**Figure 1g-i**) properties revealed a progressive maturation of intrinsic properties across the ages tested broadly in line with previous reports ^18^. However with a number of properties – membrane time constant (tau; **Figure 1f**), rheobase (pA; **Figure 1g**) and maximum firing frequency (Hz; **Figure 1i**), there was increased variability (±SD) with age (typically P5 onward) that suggests that not all SPNs acquired the same degree of maturity as development progressed.

**Figure 1.**
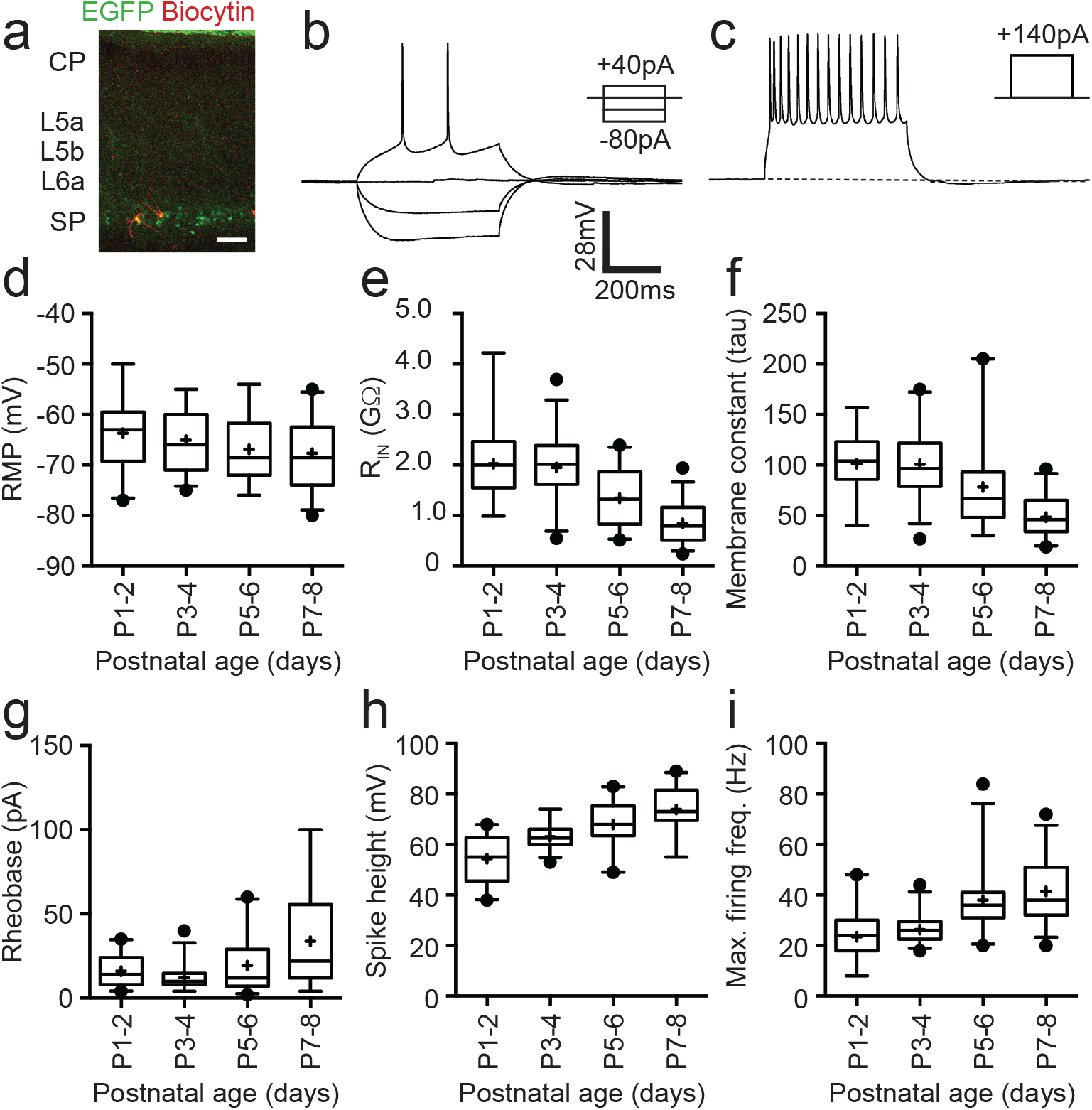
Intrinsic electrophysiological properties of *Lpar1-EGFP* SPNs. (a) Streptavidin (568nm) labelling of record *Lpar1-EGFP* SPNs in mouse S1BF at P2. (b) Superimposed electrophysiology traces recorded from one of the cells shown in panel (a) in response to hyperpolarizing and depolarizing threshold current injection. (c) Maximum firing frequency for the same cell; scale bar is the same for panels (b) and (c). (d-i) Summary data for 103 cells grouped according to age: P1-2 (n=22); P3-4 (n=27); P5-6 (n=24); P7-8 (n=30). The range of passive membrane properties recorded included (d) resting membrane potential (RMP)(mV); (e) Input resistance (R_IN_)(GΩ); (f) Membrane time constant (tau)(ms). Active properties included (g) current injection required for threshold spike (rheobase)(mV); (h) spike amplitude (mV); (i) maximum firing frequency (Hz).

Inclusion of Biocytin in the intracellular solution allowed us to reveal the morphologies of recorded SPNs (**Figure 2a**); both those assessed for intrinsic electrophysiological profile and subsequent experiments. In total we recovered 58 morphologies of 103 recorded SPNs that showed complete or near complete preservation of both axonal and dendritic arbor. It was evident from our reconstruction of 19 of these cells that Lpar1-EGFP SPNs fell into two broad categories based on dendritic arbor (**Figure 2b**): (1) fusiform SPNs with bitufted dendrites extending horizontally to the white matter tract; (2) pyramidal-like SPNs with a prominent apical dendrite projecting into layer (L)6a. Analysis of the directionality of dendritic arbor (**Figure 2c**) confirmed two distinct populations based on this criterion alone. Overlaying the axonal arbor revealed a further difference in the morphology of these two populations: the axon of pyramidal SPNs (**Figure 2d**) projected to the marginal zone/L1 with collaterals present within L6a and the subplate (SP) or layer 6b (L6b). The axon of fusiform cells was however largely restricted to the SP with a few collaterals extending into L6a (**Figure 2e**). Both cell types had extensive, but relatively simple, axonal arbors that often extended beyond the field of the low power photomicrograph either through L1 (pyramidal) or SP (fusiform). It was evident that these long-range projections extended beyond S1BF to adjacent cortical areas such as secondary somatosensory cortex (S2). Morphologies were recovered across all recorded ages, however, the proportion of fusiform cells decreased from P5 onward (**Figure 2f**). Previous analysis of the neurotransmitter phenotype of *Lpar1-EGFP* SPNs was conducted at P7 ^7^, a time point when fusiform SPNs are no longer present in our sample. To explore the possibility that these cells represent a transient GABAergic SP population^26^ we performed immunohistochemistry for GABA at P3 (**Figure 2g**). This confirmed that EGFP+ cells in the SP were all GABA-negative (n=3 brains) while profiles in L5b were GABA+ consistent with our previous characterization of the SP^7^ and L5b interneuron populations^2^. These data identify *Lpar1-EGFP* SPNs as glutamatergic projection neurons that fall into two subtypes based on their morphology: fusiform cells that innervate the SP, and pyramidal cells whose axons extend through the entire depth of the developing cortex to ramify extensively through the margin zone/L1 (**Figure 2h**). We found no evidence of selectively targeted axonal innervation of L4 by either subtype at the ages recorded. However, we cannot discount innervation of L4 glutamatergic spiny stellate neurons via their apical dendrites extending to L1, which transiently exist prior to the end of the first postnatal week^27^. Finally, these data suggest that the fusiform population of *Lpar1-EGFP* SPN is a transient population of EGFP+ SPN not present in mature cortex

**Figure 2.**
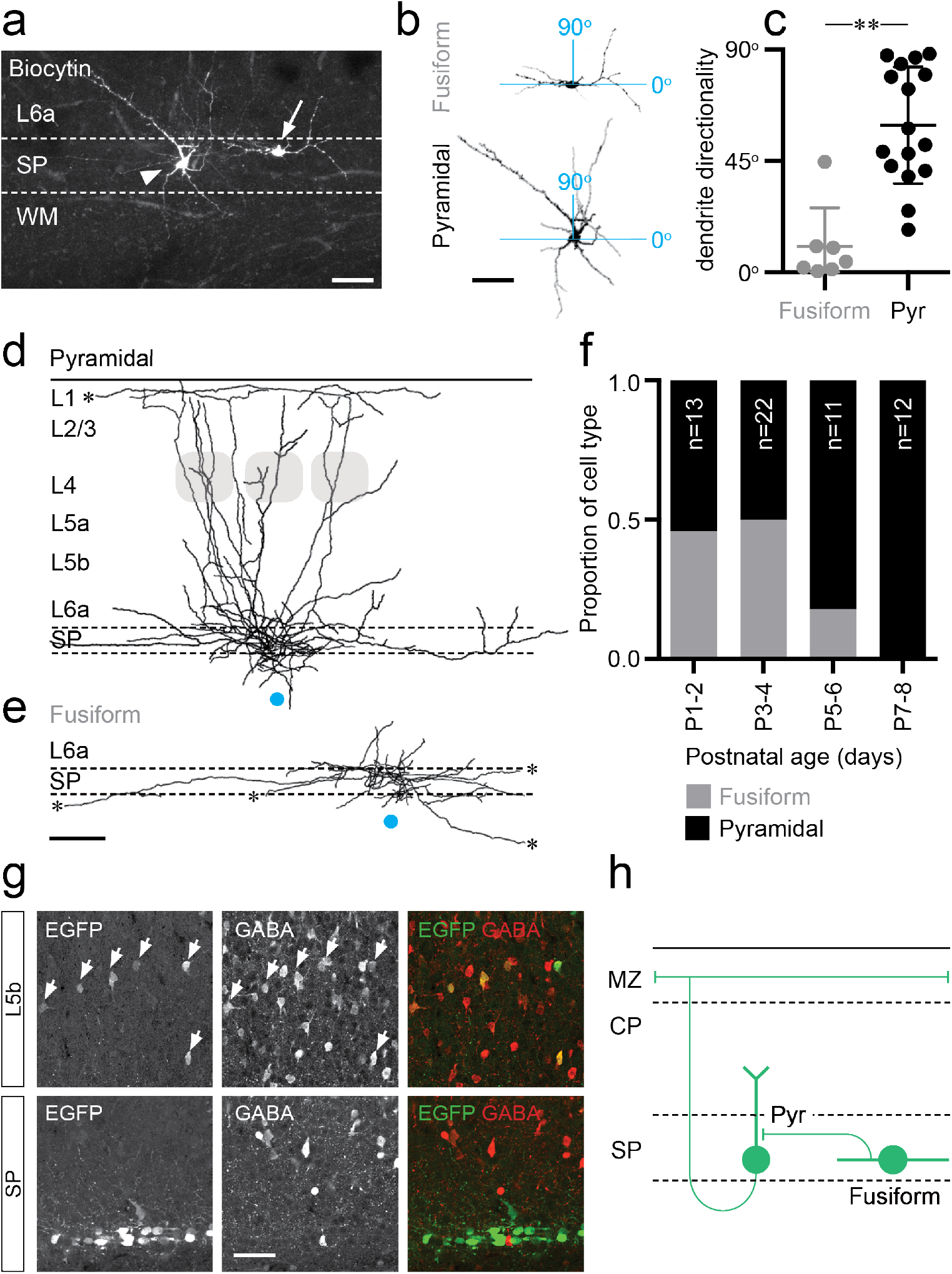
Two distinct morphologies of *Lpar1-EGFP* SPNs in S1BF. (a) Streptavidin labelled morphologies of 2 EGFP+ SPNs recorded at P3; arrowhead, pyramidal subtypes with apical dendrite projecting at ~45° into L6a; arrow, fusiform morphology with horizontal, bitufted dendrites largely restricted to the SP (Scale bar = 25μm). (b) Reconstructed (ImageJ) dendritic arbors of the cells shown (a) were used to calculate directionality with 90° indicative of vertically orientated dendrites and ~0° primarily horizontal dendrites. (c) A difference was observed in the dendritic orientation of fusiform and pyramidal SPNs (Mann-Whitney U=5, fusiform n=7, median = 4.38; pyramidal n=12, median = 54.89, **P<0.001 two-tailed). Overlay of axonal arbors of (d) pyramidal (recovered between P1-P8) and (e) fusiform (recovered P1-P5) cells aligned on soma location (horizontal position indicated by the blue circle); approximate barrel location indicated by grey shaded areas for P5+ cells; scale bar 180μm. (f) Proportion of fusiform (grey) and pyramidal (black) *Lpar1-EGFP* SPNs over the first postnatal week. (g) Immunohistochemistry for EGFP (left panel) and GABA (middle panel); EGFP+/GABA+ cells indicated by the white arrows; right panel, overlay of EGFP (green) and GABA (red). (h) Schematic of the two morphological subtypes of *Lpar1-EGFP* SPN: pyramidal (Pyr) and fusiform encountered prior to P5. MZ, marginal zone; CP, cortical plate; SP, suplate.

### Laser scanning photostimulation reveals dynamic synaptic integration of *Lpar1-EGFP* SPNs into local glutamatergic network

We used UV (355nm) laser photolysis of caged glutamate to map afferent synaptic input onto SPNs in acute *in vitro* cortical slices (**Figure 3a**). We mapped input using laser scanning photostimulation (LSPS) across the extent of a pseudo-random (50μm spaced) grid covering the depth of neocortex immediately above any given recorded cell. From the earliest time points recorded (P1-2) SPNs received distinct columnar glutamatergic input, either from SP and adjacent cortex (L6a) alone, or from both SP/L6a and more superficial cortex; patterns of innervation that we termed ‘local’ and ‘translaminar’ respectively (**Figure 3b**). The average laminar profile of local (n=15) and translaminar (n=8) SPNs at P1-2 revealed that the latter received input from the cortical plate (CP) absent in local SPNs (**Figure 3c**). This translaminar input became more prominent over the next two postnatal days (P3-4; **Figure 3d**) with the source primarily focused in the lower CP, presumptive L4. In contrast, at P5-6 we recorded relatively few translaminar neurons with the majority (84%) dominated by local SP/L6a input. The 3 cells defined as translaminar received afferent input from mostly infragranular pyramidal cells (**Figure 3e**). This trend continued in the cells recorded at P7-8 although the latter were diverse in input source such that the average laminar profile and map of translaminar cells resembled a diffuse columnar band of glutamatergic input (**Figure 3f**) across L4 and L5b.

**Figure 3.**
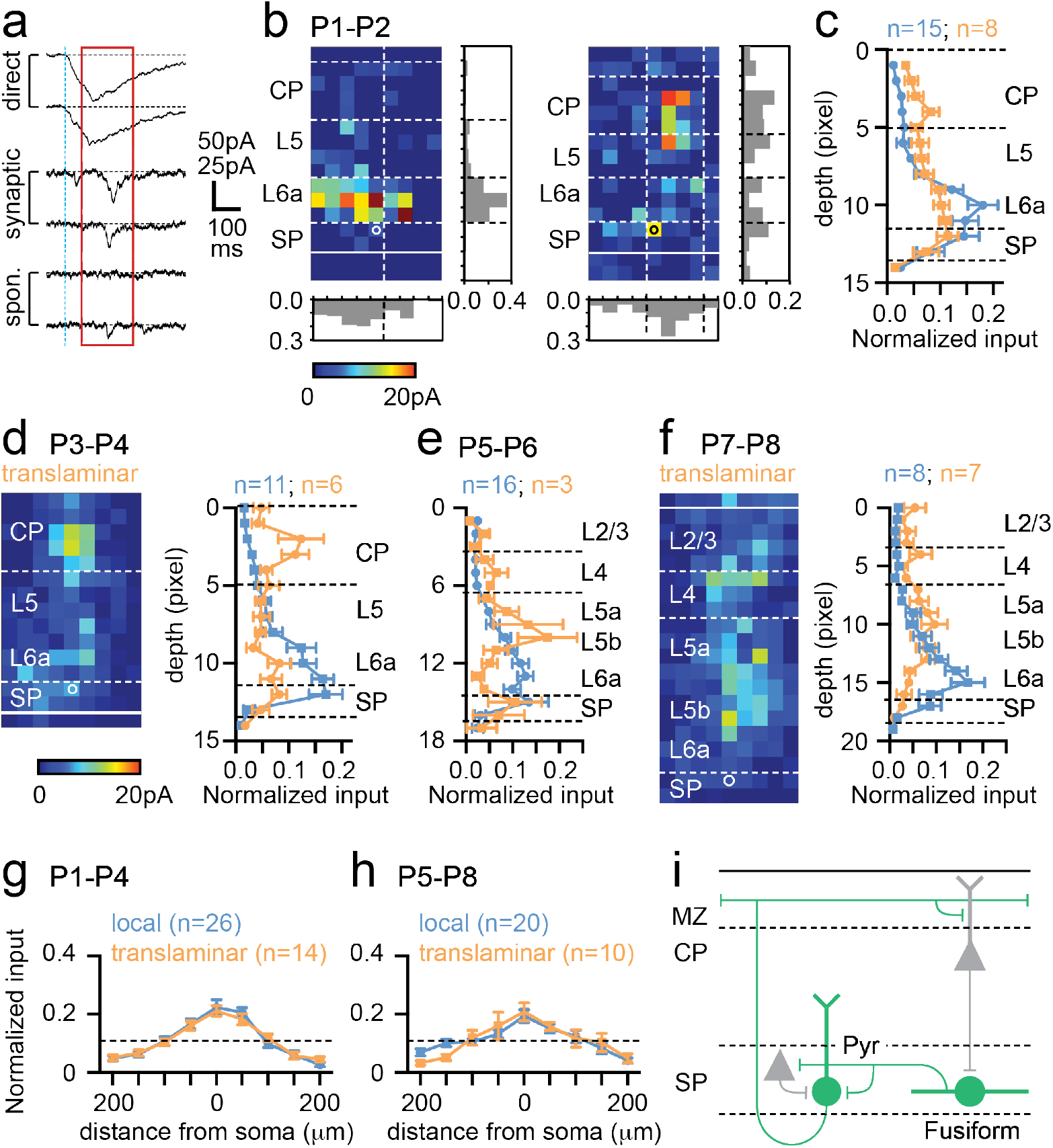
Synaptic integration of *Lpar1-EGFP* SPNs into the local cortical glutamatergic network. (a) LSPS of caged glutamate resulted in 3 responses observed in whole cell patch clamp recordings of SPNs: top traces, large amplitude direct responses with onset locked to laser pulse onset (dashed vertical blue line); middle traces, synaptic response of consistent EPSCs within the monosynaptic event window (red box); bottom traces, no consistent response with occasional spontaneous EPSCs. Scale bar for direct traces: 50pA; for synaptic and spontaneous: 25pA. (b) Local (left) and translaminar (right) glutamatergic input maps for SPNs recorded at P1-2. Pixel size: 50μm. (c) Average input profile for local (blue) and translaminar (orange) SPNs; translaminar SPNs showed increased input from the cortical plate (CP) and reduced local (L6a/SP) innervation. (d) Left panel, average input map for translaminar SPNs recorded P3-P4 (n=6); right panel, average input profile for local (blue) and translaminar (orange) SPNs. (e) corresponding input profile for P5-P6. (f) Average input map and profile for SPNs recorded between P7-P8. Horizontal profile for local and translaminar SPNs aligned on cell soma at (g) P1-P4 and (h) P5-P8. (i) Schematic showing glutamatergic circuit onto *Lpar1-EGFP* SPNs.

While there were clear differences in laminar input profile, the average horizontal profile for both local and translaminar SPNs did not vary through development (**Figure 3g,h**). Recovered morphologies identified SPNs that received translaminar input from the cortical plate, presumptive L4, as being the fusiform subtype (5/6 fusiform SPNs were translaminar), whereas pyramidal SPNs received local input (5/5 SPNs). Only one fusiform cell was recovered from P5 onwards and along with the majority of pyramidal cells (6/9) this cell received local glutamatergic input. The remaining 3 pyramidal SPNs received translaminar input from L5. Taken together these data suggest that transient fusiform SPNs are the primary recipients of early translaminar input from the cortical plate (**Figure 3i**), up until P4 when their numbers start to decline. In parallel, pyramidal SPNs are dominated by local glutamatergic input from SP/L6a at early ages (**Figure 3i**), but acquire varied translaminar input from more superficial layers from P5 onward.

### Lpar1-EGFP SPNs receive distinct sources of GABAergic input including translaminar input from SST+ interneurons

Our intrinsic electrophysiology, morphological and LSPS data supports the notion that there is a reconfiguration in the *Lpar1-EGFP* SPN circuit around P5 in line with a previous report of a shift to columnar network activity in S1BF at this time^25^. To understand if this also involves early GABAergic circuits, we repeated our LSPS experimental strategy with the cell voltage clamped at the reversal potential for glutamate (E_Glut_)(**Figure 4a,b**). SPNs were pooled into two groups based on age: SPNs recorded prior to the P5 transition (P1-4; **Figure 4c-e**) and P5 onward (P5-8; **Figure 4f-h**). Similar to our previous assessment of glutamatergic input, it was evident that SPNs received either local (**Figure 4c,f**) or translaminar input (**Figure 4d,g**) across both time windows. Prior to P5, SPNs with local (**Figure 4c**) and translaminar (**Figure 4d**) input were evident in similar numbers with the latter receiving prominent columnar input from L5 (**Figure 4b,d,e**). Local GABAergic input was distributed through SP and adjacent L6a (**Figure 4c**). The average translaminar input (**Figure 4e**) revealed largely complementary distributions in GABAergic input for these two populations. From P5 onward we observed primarily local GABAergic synaptic input onto SPNs (n=11/16)(**Figure 4f**) with the source of translaminar GABAergic input highly variable in location resulting in a diffuse average input profile (**Figure 4g**) with a more-or-less even distribution across the depth of cortex (**Figure 4h**). Unlike glutamatergic input, the horizontal or columnar input was evenly spread for GABAergic input with the exception of translaminar input from P1-P4 (**Figure 4i,j**). Indeed translaminar GABAergic input onto SPNs prior to the emergence of whisker barrels at ~P5 was highly focused within the immediate column (**Figure 4i**). Lpar1-EGFP SPNs that received such input were largely fusiform (n=3 out of 4 recovered morphologies) whereas SPNs that received broadly distributed, local GABAergic input were primarily pyramidal subtypes (**Figure 4k**) as were SPN morphologies for both local and translaminar GABAergic input at P5-P8 (n=4 recovered morphologies).

**Figure 4.**
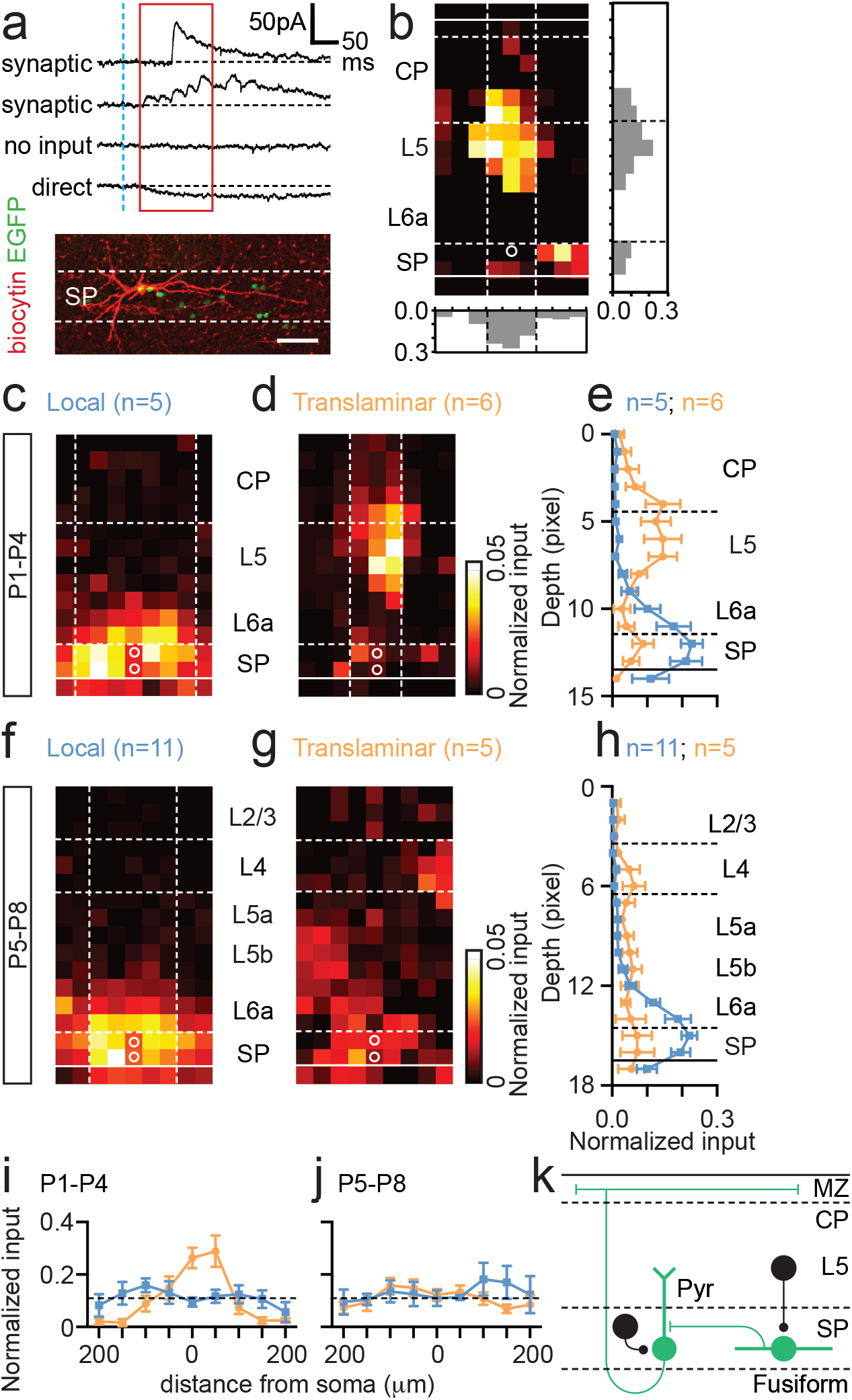
GABAergic input onto Lpar1-EGFP SPNs in the first postnatal week. a) Top panel, synaptic and small inward direct response observed in whole cell patch clamp recordings of SPNs voltage clamped at the approximate reversal potential for glutamate (E_Glut_). Bottom panel, recovered fusiform morphology of the SPN with translaminar GABAergic input map shown in (b). (c-e) GABAergic input onto SPNs recorded from P1-P4. Average input maps for (c) local and (d) translaminar SPNs with profile shown in (e). (f-h) corresponding data for SPNs recorded between P5-P8. (i,j) Columnar analysis of GABAergic input on SPNs at (i) P1-P4 and (j) P5-P8. (k) Schematic of GABAergic input onto fusiform and pyramidal (Pyr) SPNs present from P1-P4

Somatostatin (SST+) interneurons form a key component of early postnatal translaminar circuits^2, 28^ and have been shown to drive synapse and circuit maturation^29, 30^. To test if these interneurons influence SPNs and the circuit transition observed around P5 we first conditionally expressed Channelrhodopsin2 (ChR2) in SST+ interneurons by crossing mice homozygous for the *Ai32* (ChR2) reporter allele onto our *Lpar1-EGFP* background that was also homozygous for the *SST-Cre* driver line to generate *Lpar1-EGFP;SSTCre;Ai32* offspring. We then used wide field blue light (470nm) illumination to evoke SST+ interneuron IPSCs in EGFP+ SPNs voltage clamped at E_Glut_ at P3-4 (n=7) and P5-6 (n=5). We observed an increase in IPSC amplitude across all laser powers tested greater or equal to minimal stimulation in the P5-6 when compared to the P3-4 time window (**Figure 5a**). To test if the increase in amplitude was a result of either increased quantal size or number of innervations we repeated these experiments in ACSF in which extracellular Ca^2+^ was replaced with strontium (Sr^2+^). Incubation with ACSF containing Sr^2+^ (Sr-ACSF) leads to asynchronous vesicular release at the presynaptic terminal providing a reasonable estimate of quantal size^31, 32^. In our hands incubation of neonatal SPNs in Sr-ACSF resulted in asynchronous release observed at minimal stimulation (**Figure 5b**) and a significant difference in ChR2-dependent IPSC amplitude between control and Sr-ACSF conditions (**Figure 5c**). However we observed no difference in the amplitude of IPSCs recorded in Sr-ACSF between P3-4 and P5-6 time windows (**Figure 5d**) despite the significant difference in amplitude under control conditions (**Figure 5b,c**). This suggests the observed increase in amplitude at this time point results from an increase in innervation by SST+ interneurons rather than an increase in quantal size.

**Figure 5.**
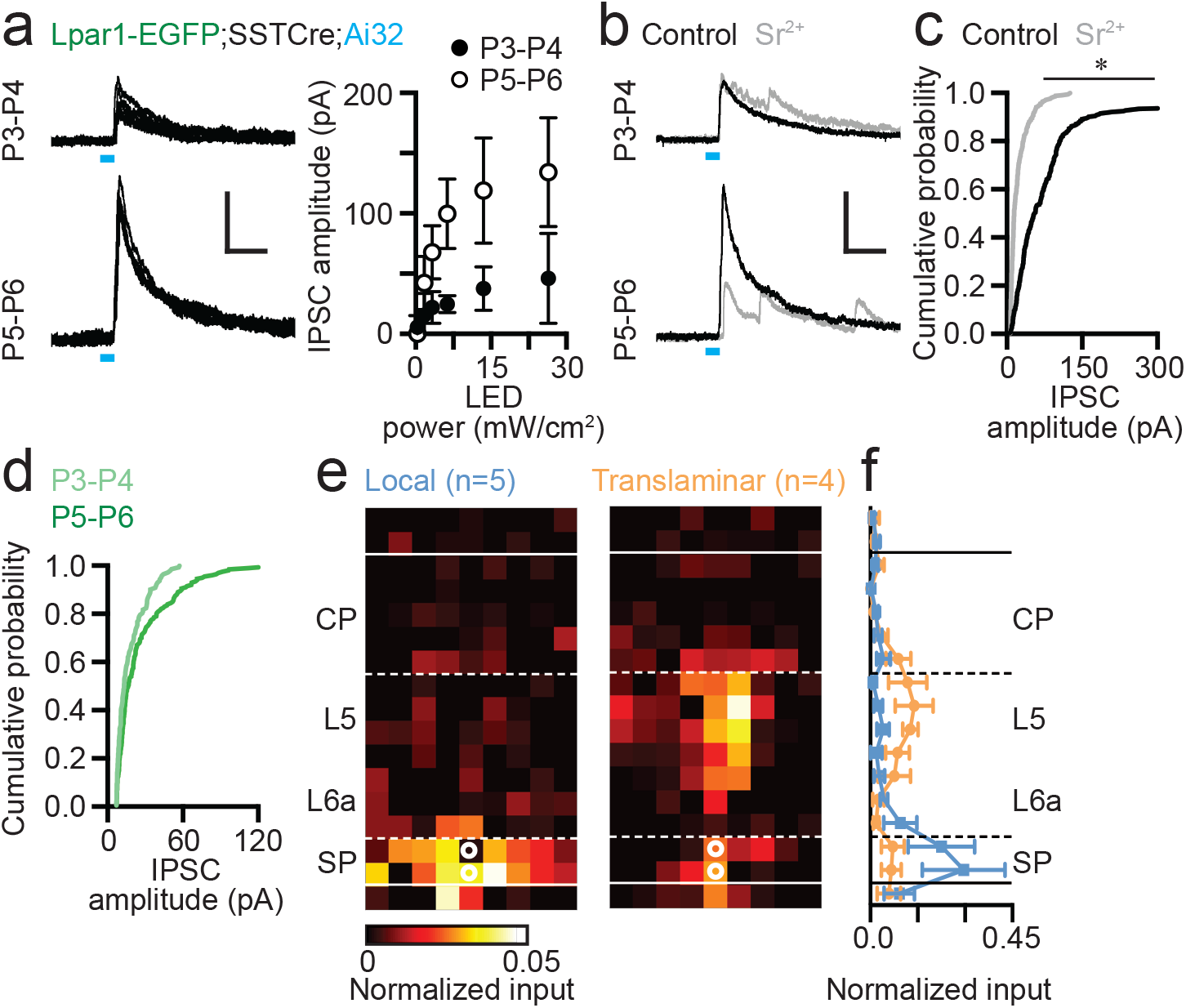
Somatostatin-positive (SST+) interneurons innervate *Lpar1-EGFP* SPNs during early postnatal life. (a) Widefield 473nm blue light stimulation evoked IPSCs in SPNs at both P3-P4 and P5-P6 following conditional expression of Channelrhodopsin2 (ChR2) in SST+ interneurons. LED on indicated by the blue line. (b) Incubation in Sr^2+^-containing ACSF resulted in asynchronous neurotransmitter release. (c) Cumulative probability plot of ChR2-evoked IPSC amplitude for control (black line) versus Sr^2+^-containing HDC ACSF (grey) across P3-P6; asterisk: two-sample Kolmogorov-Smirnov test: P ≤ 0.01. (d) Comparison of early (P3-P4) versus late (P5-P6) IPSC amplitude in the presence of Sr^2+^-containing HDC ACSF. (e) Local and translaminar SST+ interneuron input maps onto SPNs revealed through LSPS uncaging of ATP in conjunction with conditional expression of P2X2 receptor in SST+ interneurons. (f) Local (blue) and translaminar (orange) average layer profiles for SST+ input onto SPNs.

Having established that SPNs received SST+ interneuron input through the first postnatal week, we next employed conditional expression of the P2X2 receptor – an optogenetic actuator that we have previously used in conjunction with uncaging of ATP^28^, that allows us to map the somatic location of presynaptic SST+ interneurons. LSPS uncaging of ATP over the 50μm spaced pseudorandom grid revealed two distinct input profiles for SPNs in the immediate postnatal window (P1-P4)(**Figure 5e**): local SP/L6a *versus* translaminar infragranular SST+ interneuron input. Similar to our previous findings with both glutamatergic and global GABAergic input, local input SPNs were pyramidal cells (3/3 recovered morphologies) whereas SPNs that received translaminar synaptic input were predominantly fusiform (2/3). At the later time point (P5-8) SST+ interneuron input onto SPNs was exclusively local (n=6)(*data not shown*); an observation that fits with previous data supporting the presence of a transient network between superficial cortical layers and fusiform SPNs. The presence of GABAergic input onto SPNs from infragranular SST+ interneurons precedes our previously reported reciprocal connection between these SST+ cells and L4 spiny stellate neurons during the L4 critical period for plasticity^2^ and then onto L2/3 pyramidal cells during the emergence of L4 to L2/3 feed-forward connections^28, 33^. Taken together, this evidence suggests that infragranular SST+ interneurons sequentially innervate thalamo-recipient layers through early postnatal life in S1BF.

### Sparse thalamocortical input onto early postnatal *Lpar1-EGFP* SPNs in S1BF

SPNs are thought to play an important role in early thalamic integration in primary sensory cortices. To assess the role that *Lpar1-EGFP* SPNs play in the early sensory circuit we first used electrical stimulation of the ventrobasal complex (VB) of the thalamus while recording from EGFP+ SPNs in acute *in vitro* thalamocortical slice preparation (**Figure 6a**). Electrical stimulation evoked excitatory postsynaptic currents (EPSCs) in the majority (76%) of SPNs recorded across the time window studied although there was a drop in incidence between the P3-4 (92%; n=12 cells) and P5-6 (61%; n=18) time window (**Figure 6b**); an absence of thalamic input onto any given SPNs was only recorded if TC-EPSCs were observed in other SPNs or layer 4 neurons in the same thalamocortical slice. Analysis of the amplitude of the minimal electrical stimulation EPSC (**Figure 6c**) identified a significant change in variance between these times but no difference in amplitude (P = 0.08; two-tailed t-test, t=1.92, df=11.19). To further identify putative thalamocortical EPSCs (TC-EPSCs) we recorded the latency of the evoked EPSC, jitter (standard deviation in latency, ms), and amplitude for the minimal stimulation EPSC for 53 SPNs. All EPSCs with a latency >10ms and/or jitter >1.0ms (**Figure 6d**) were then excluded from our analysis leaving 32 SPNs that could be further divided into two populations based on 10-90% rise time and amplitude (**Figure 6e**): type 1 (n=9), large amplitude (55.3 pA ± SD 14.3), low jitter (0.20 ms ± SD 0.07), *versus* type 2 (n=23), small amplitude (14.9 mV ± SD 5.8), high jitter (0.41 ms ± SD 0.17) EPSCs (**Figure 6f**). Whether both populations represent TC-EPSCs^32, 34^ was unclear from our electrical stimulation of VB in part because both conform to a previous criteria used to distinguish TC-EPSCs from antidromic cortico-thalamic EPSCs namely that TC-EPSCs exhibit standard deviation in jitter <1.0ms ^35^. However thalamic connectivity could be as low as 17% if type 1 EPSCs recorded in SPNs represent true orthodromic TC-EPSCs; considerably lower than connectivity reported in previous studies of SP in early postnatal ages.

**Figure 6.**
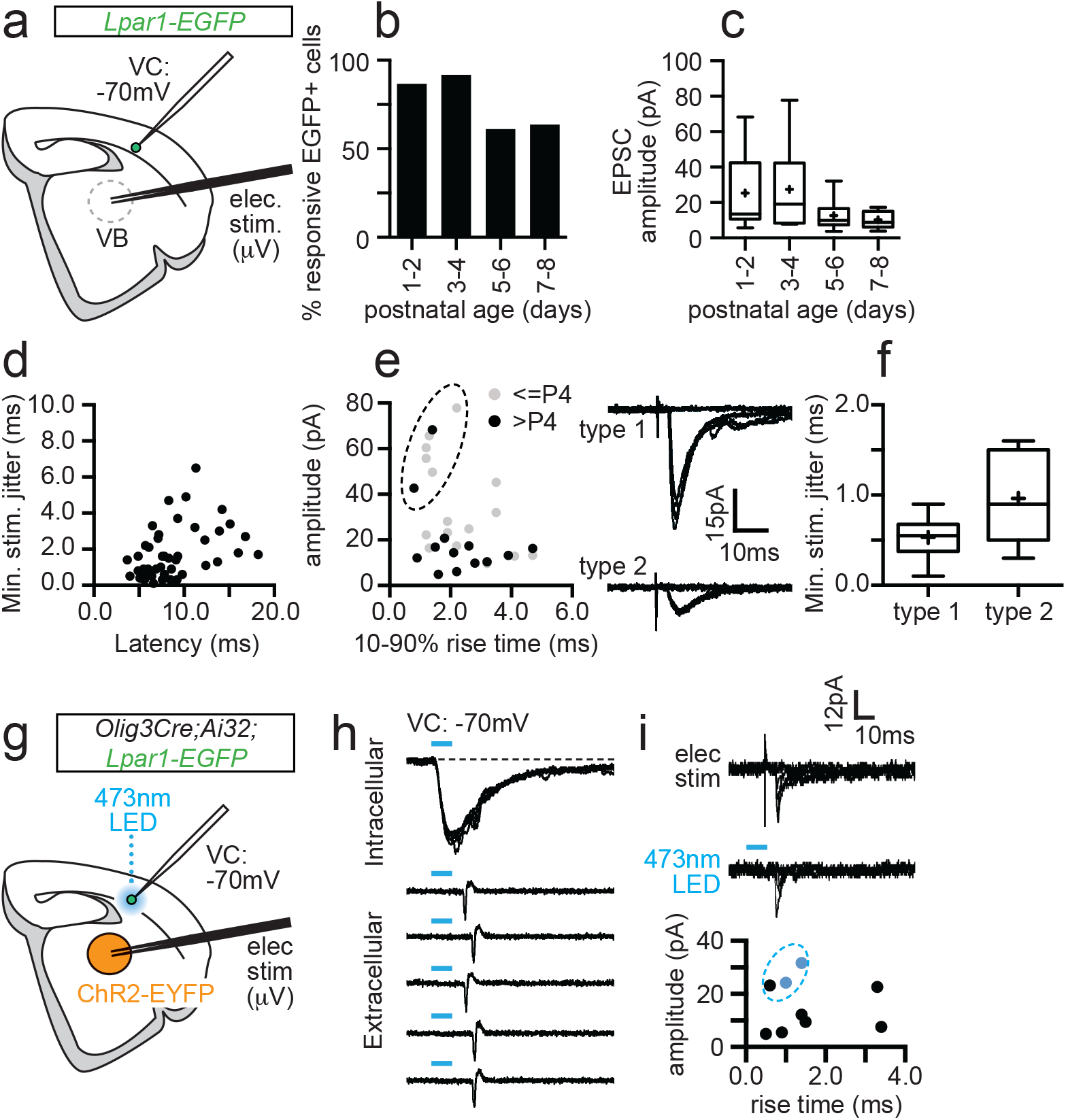
Sparse thalamic afferent input onto *Lpar1-EGFP* SPNs in postnatal S1BF. (a) Schematic showing the experimental set-up for recording electrical stimulation-evoked TC-EPSCs in thalamocortical slices. (b) Percentage EGFP+ SPNs exhibiting constant latency synaptic response to minimal electrical stimulation over development. (c) Box plot showing the average amplitude for EPSCs. (d) Plot of jitter (Standard deviation of onset EPSC) versus average latency for all the responses shown in (b,c). Putative TC-ESPCs had latency ≤ 10ms and jitter ≤ 1ms. (e) Plot of average EPSC amplitude (pA) versus 10-90% rise time for all putative TC-EPSCs. Dashed circle, type 1 TC-EPSCs (top, right trace) versus type 2 (bottom trace). (f) Box plot of average jitter for type 1 and type 2 EPSCs. (g) Schematic showing the experimental set-up for combined electrical and optogenetic stimulation of thalamic afferents. (h) (top panel) Blue light reliably evoked inward currents in thalamic relay neurons following *Olig3 Cre* conditional expression of ChR2, sufficient to trigger action potentials in loose cell attached recordings (bottom panel). (i) Electrical (top trace) and optogenetic (bottom trace) stimulation evoked TC-EPSC in a single *Lpar1-EGFP* SPN at P2. Bottom panel, plot of electrical stimulation evoked TC-EPSCs recorded during the combined electrical and optogenetic stimulation experiments. Blue data points indicate the two cells that also exhibited optogenetic EPSCs; dashed blue line, TC-EPSCs with a type 1 profile.

Given the disparity with previous reports we decided to employ optogenetics in parallel with electrical simulation of VB to unequivocally identify EPSCs arising from thalamic input (**Figure 6g**). We conditionally expressed ChR2 in thalamic nuclei using the *Olig3* Cre driver line that causes recombination throughout the thalamus early in development^36, 37^. To validate our optogenetic strategy at early postnatal ages, we first recorded from thalamic relay neurons in *Lpar1-EGFP;Olig3 Cre;Ai32* mice and established that we could evoke (1) reliable inward currents in whole cell patch clamp mode in response to blue, 470nm light (**Figure 6h**, top panel), and (2) time-locked action potentials in cell-attached mode (**Figure 6h**, bottom panel) from the earliest time points recorded (P1)(n=5 VB cells). We then recorded *Lpar1-EGFP* SPNs and tested for TC-EPSCs using both electrical and light stimulation protocols. Across the whole time window tested (P1-8) we obtained recordings under both stimulation protocols in 19 EGFP+ cells. Of these, 9 cells across all ages tested (P1-8) had short latency, low jitter (<1ms) EPSCs in response to electrical stimulation (type 1 and 2)(**Figure 6i)**. However only 2 cells (11% of *Lpar1-EGFP* SPNs), with properties consistent with type 1 EPSCs, exhibited responses to both 470nm light and electrical stimulation; no EGFP+ cells showed synaptic responses to light alone. One cell was recorded at P2 and was fusiform in morphology; the second was a P5 pyramidal SPN. To confirm our ability to resolve thalamic input using this dual stimulation strategy we further recorded non-EGFP+ SPNs and L4 cells in the earlier time window (<P5) and found 40% (n=5) and 33% (n=3) connectivity respectively. Taken together these data suggest that *Lpar1-EGFP* SPNs in S1BF receive thalamic input in the first postnatal week but that this input is relatively sparse, contacting only a small subset of the total population.

## DISCUSSION

We have recorded from a genetically identified population of subplate neuron (SPN) through the first postnatal week to establish the contribution of this cell type to early circuits of somatosensory whisker barrel cortex (S1BF). Our data reveal that the *Lpar1-EGFP* population represents two distinct subtypes with different somatodendritic morphologies and with afferent input throughout the first 4 postnatal days (P1-4): first, pyramidal SPNs that receive primarily local input at this stage from the subplate (SP) network, but whose axons traverse across all the layers of cortex to project horizontally via layer 1 (**Figure 7a**). Second, fusiform SPNs that receive translaminar input from more superficial cortical layers but whose axons are largely confined to the immediate layer and therefore output to the SP network (**Figure 7b**). Fusiform SPNs receive glutamatergic input from the cortical plate, including putative layer 4, as well as GABAergic input from infragranular SST+ interneurons; two signalling centers that form reciprocal connections through the L4 critical period of plasticity in S1BF^2^. During the later period (P5-8) fusiform cells are encountered in significantly reduced numbers – indeed are absent in the P7-8 window. The remaining pyramidal SPNs are evenly split between those that are still dominated by local inputs and those that acquire an array of diverse inputs from across cortical layers, located both in the immediate and adjacent cortical columns. This diversification of synaptic input fits with the rapid transition to columnar signalling previously reported at P5 ^25^, and suggests that this time point represents the switch from transient SP to layer 6b (L6b) ^13^ in S1BF.

**Figure 7.**
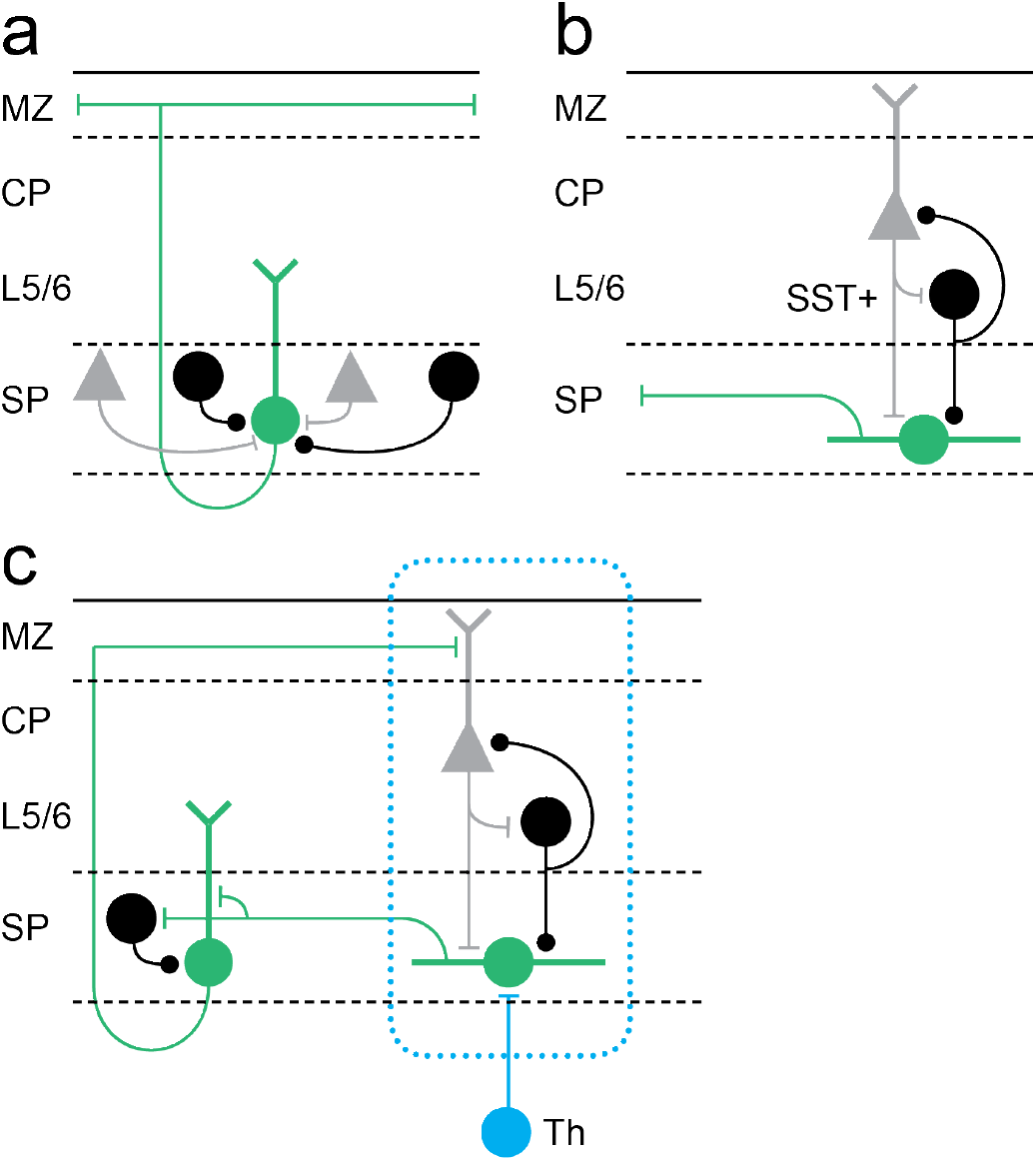
Model for *Lpar1-EGFP* SPN circuits in early postnatal cortex. (a) Pyramidal *Lpar1-EGFP* SPNs received broad glutamatergic (grey neurons) and GABAergic (black) input from the subplate zone. (b) Transient fusiform *Lpar1-EGFP* SPNs in contrast receive translaminar input from glutamatergic neurons (grey) in the cortical plate and SST+ interneurons (black) in infragranular layers. (c) We propose that sparse thalamic input (Th) onto *Lpar1-EGFP* SPNs interacts with both fusiform and pyramidal SPN circuits to sculpt the emergent columnar cytoarchitecture (blue dashed box).

Our study identifies that *Lpar1-EGFP* SPNs in S1BF have an additional novel function that does not conform to the canonical model for SPNs established across sensory cortices, wherein SPNs act as relay cells for thalamic input to L4 ^1, 4, 38^. We have tested thalamic engagement with Lpar1-EGFP SPNs using combined electrical and optogenetic stimulation of thalamic afferent fibres and found that the incidence of connectivity onto postnatal *Lpar1-EGFP* SPNs to be as low as 11% in thalamocortical slices that otherwise showed good preservation of connectivity. This is at odds with a number of previous studies reliant exclusively on electrical stimulation^18, 39^ which obtained levels of connectivity approaching our initial electrical stimulation paradigm. While this could be a property of the *Lpar1-EGFP* component of the subplate, it is definitely worth revisiting perinatal thalamic engagement with the cortex using optogenetic approaches given that conditional expression of ChR2 in thalamic nuclei allows unequivocal discrimination of thalamocortical versus corticothalamic input without possible antidromic activation. Differences in the level of thalamic engagement aside, our data are consistent with the model that thalamic input is amplified via the subplate network^40^ and onward communicated via pyramidal SPNs to more superficial layers of cortex via layer 1. None of our recovered morphologies suggest dense innervation of L4 *per s*e by this particular genetically-defined subpopulation of SPN^39^. Indeed this and our extensive mapping of S1BF using LSPS through early postnatal life^28, 41^ provide little evidence that there is a privileged route of connectivity between SP and L4 for a protracted period during early postnatal life in this primary sensory area, in contrast to other sensory modalities.

We targeted a genetically-defined population of SPN using the *Lpar1-EGFP* transgenic mouse line (GENSAT). Lpar1 (Edg2) is one of a number of markers previously shown to delineate SPN diversity^7^ with the cohort labelled by this line one of the earliest born subtypes with peak neurogenesis at embryonic day (E)11.5. EGFP expression is evident in this population at embryonic ages^7^, increasing to label both SP and non-SP neurons by the end of the first postnatal week^2^. This reported increase in strength of EGFP expression in SPNs through early postnatal life would seem to preclude the possibility that the fusiform SPN subtype down-regulate EGFP at P5. The most parsimonious explanation – that reconciles our observation of fusiform *Lpar1-EGFP* cells as a ‘classical’ transient SPN population with the fact that there is no significant decrease in the number of EGFP+ SPNs over this time window^7^ – is that this subtype represent only a small fraction of the EGFP+ SPN number. Our observed increase in the prevalence of the pyramidal subtype of EGFP+ SPNs over development has been documented by others^12^. Finally the *Lpar1-EGFP* transgenic is an excellent tool for targeting and recording these two morphological variants of SPN. However, our knowledge of the efferent targets of these cells is, in absence of a conditional genetic strategy, limited to a purely morphological assessment. Alternative genetic approaches such as CRE-DOG^42^ although useful to researchers targeting EGFP+ neuronal populations in the adult cortex^43^ do not provide a viable means of targeting the *Lpar1-EGFP* cells within the first postnatal week. What is evident from our analysis is that the two subtypes present in the earlier time window target completely different layers within the developing cortical plate, highly suggestive of different roles within the early circuitry of S1BF.

Laser scanning photostimulation (LSPS) has been used previously in conjunction with glutamate uncaging to probe the early SP circuits of primary auditory cortex (A1)^10, 15^. Similar to these reports, we find SPNs that receive local and translaminar input. However the temporal dynamics of the connections that we observe in S1BF, notably the early afferent input from L4, are quite different to those reported for A1 wherein L4 input emerges in the second postnatal week^10^. This disparity in timing could underpin differences in the role of SP in circuit maturation between sensory areas and suggests that the cytoarchitecture of sensory areas differs from the onset.

In recent years optical approaches have extended our knowledge of the early circuit of somatosensory cortex. It is evident that prenatal spontaneous thalamic activity plays an instrumental role in determining the columnar organisation of S1BF^24^. Shortly before birth thalamic afferent fibres are restricted to the subplate and thalamic stimulation elicits activity that spreads laterally through this and immediate adjacent infragranular layers^17^. It is possible that at these early ages thalamic innervation of SPNs is widespread in S1BF and that the role of such spontaneous activity is to competitively select the sparse SPNs that will maintain thalamic innervation into the first few postnatal days; a time point by which sensory activity has already transitioned to the overlying cortical layers in S1BF^24^. Our data suggest that *Lpar1-EGFP* pyramidal SPNs are likely conduits for such activity, recruiting pyramidal cells and GABAergic interneurons in more superficial cortex^14, 44^ via their L1 axon collaterals. In turn, neurons in the superficial cortical layers provide feedback columnar glutamatergic and GABAergic synaptic input onto transient fusiform SPNs thus completing the circuit (**Figure 7c**). This indirect mechanism could provide the necessary framework for the interpretation of early thalamic signals resulting in columnar organization. Indeed, such a mechanism coupled with the sparse nature of thalamic engagement with the subplate, represents a plausible substrate to ensure the emergence of spatially distinct columnar circuits in S1BF. Moreover, relay of thalamic input via L1 moves away from a L4-centric view of early thalamic engagement, more in line with distributed thalamocortical input across all 6 layers of neocortex^45^. Our findings provide insight into the earliest neuronal networks of somatosensory cortex, highlighting transient glutamatergic and GABAergic circuits that are essential for the emergence of normal perception.

## METHODS

### Animal husbandry and use

Animal care and experimental procedures were approved by the University of Oxford local ethical review committee and conducted in accordance with UK Home Office personal and project (70/6767; 30/3052; P861F9BB75) licenses under the Animals (Scientific Procedures) 1986 Act. The following mouse lines were used: *Lpar1-EGFP* (Tg(Lpar1-EGFP)GX193Gsat), *SST-ires-Cre* (Sst^tm2.1(cre)Zjh^/J), *Olig3 Cre* (Olig3^tm1(cre)Ynka^), *Ai32* (RCL-ChR2(H134R)/EYFP), *R26::P2X2R-EGFP* (floxed-stop-rat P2X2 receptor). All experiments were performed blind to the mouse genotype with the exception of *Lpar1-EGFP* transgene which is Y chromosome-linked^7^. The date of birth was designated postnatal day (P)0.

### Acute *in vitro* slice preparation

Acute brain slices were prepared as previously described^2^. Male mice (postnatal day 1 to 8; P1-P8) were anaesthetised with 4% isoflurane in 100% O_2_ and decapitated; the cerebral cortex was quickly dissected in ice-cold, oxygenated (95% O_2_ / 5% CO_2_) artificial cerebrospinal fluid (ACSF) of the following composition (in mM): 125 NaCl, 2.5 KCl, 25 NaHCO_3_, 1.25 NaH_2_PO_4_, 1 MgCl_2_, 2 CaCl_2_, 20 glucose (300-310 mOsm; all chemicals were purchased from Sigma unless otherwise stated). Coronal and thamalocortical slices (350-400 μm) including the primary somatosensory barrel cortex (S1BF) were cut in ice-cold ACSF through a vibratome (Vibratome 3000 Plus; The Vibratome Company) and allowed to recover in ACSF at room temperature (RT) for at least 1 h prior to electrophysiological recordings. Coronal slices were obtained by cutting the brain at an angle perpendicular to S1BF; thalamocortical slices were obtained according to established procedures with the angle varied according to developmental age^2, 46^.

### Whole-cell patch-clamp electrophysiology

Slices containing S1BF were selected for electrophysiology experiments if they showed good preservation of the radial structure, as assessed by the presence of layer (L)5 pyramidal neuron apical dendrites extending to supragranular layers. *Lpar1-EGFP* subplate neurons (SPNs) were readily distinguished from *Lpar1-EGFP* GABAergic interneurons based on their localization in a thin layer of cells located between the cortical L6 and the underlying white matter, identified as the subplate (SP). Layer 6b could be detected as a thin, compact cell layer that could be distinguished from Layer 6a and white matter. Cells were selected ~50 μm below the slice surface and targeted for patch-clamp recordings guided through infrared-differential interference contrast (IR-DIC) microscopy using a 40x water-immersion objective. Whole-cell patch clamp electrophysiological recordings were performed at RT using a Multiclamp 700B amplifier and Digidata 1440A digitizer (Molecular Devices, USA). Patch pipettes were obtained from borosilicate glass microelectrodes (6-9 MΩ; Harvard Apparatus, UK), pulled through a PC-10 puller (Narishige, Japan). Electrodes were filled with either a K-based (128 mM K-gluconate, 4 mM NaCl, 0.3 mM Li-GTP, 5 mM Mg-ATP, 0.1 mM CaCl_2_, 10 mM HEPES; pH 7.2 with KOH; 280-290 mOsm) or Cs-based intracellular solution (100 mM gluconic acid, 0.2 mM EGTA, 5 mM MgCl_2_ 40 mM HEPES, 2 mM Mg-ATP, 0.3 mM Li-GTP; pH 7.2 with CsOH; 280-290 mOsm). Biocytin (0.3%) was included in the intracellular solution to allow the morphological reconstruction of recorded neurons. To study excitatory postsynaptic currents (EPSCs), SPNs were held at a holding potential (V_h_) of −60 mV; inhibitory postsynaptic currents (IPSCs) were recorded by voltage-clamping the cell near the equilibrium potential for glutamate (E_Glut_). During glutamate uncaging experiments, E_Glut_ was found empirically by uncaging glutamate in the proximity of the recorded cell and tuning the V_h_ until no net laser-induced direct postsynaptic current was observed; otherwise V_h_ was set to 0 mV (corrected for calculated liquid junction potential of ~13 mV for the Cs-based intracellular solution). All recordings were sampled at 20 kHz and low-pass filtered online at 0.5 kHz. Cell input and series resistance (R_in_ and R_s_) were monitored throughout the duration of the recording without applying compensation; recordings were discarded when R_s_ exceeded 20% of its initial value.

Cells recorded with the K-based intracellular solution were initially recorded in current-clamp configuration to record their intrinsic electrophysiological profile using both depolarizing and hyperpolarizing current steps (500ms-long).

### Laser-scanning photostimulation: methods and analysis

Laser-scanning photostimulation (LSPS) was performed according to the method previously described^41, 47^. This optical technique allows to stimulate neurons into a small portion of cortical tissue (~50μm) while recording the postsynaptic current in the target neurons. Thus, the location of any presynaptic neurons showing functional connectivity to the recorded one can be inferred by the location of the optical stimulation. Prior to LSPS, slices were incubated for a minimum of 6 mins. with high-divalent cation (HDC) ACSF of similar composition to the normal ACSF but with increased concentration (4 mM) of MgCl_2_ and CaCl_2_ and supplemented with 100μM MNI-caged glutamate (Tocris Bioscience, UK) for glutamate uncaging experiments. Mapping of cell-type-selective inputs were performed with an optogenetic strategy previously described^28^. In brief, the P2X2 receptor was conditionally expressed into SST+ interneurons and selectively stimulated by laser uncaging of DMNPE-caged ATP (100μM, Life Technologies, UK). LSPS was performed using an ultraviolet (UV) laser (DSPL-355/30) and a galvanometer targeting system (UGA-42, Rapp Optoelectronic GmbH, Germany) focused through a 10x Olympus objective. The stimulation grid was organized into 17×9 target spots (~50μm spatial resolution). Long-duration (100 ms), low-power (<2 mW at sample plane) laser pulses were fired in a pseudo-random order at 1-2 Hz frequency. In order to cover the whole extent of the cortical column, 2-3 LSPS grids were sequentially employed and properly aligned and averaged offline during data analysis. For each individual LSPS grid, a minimum of 3 runs were obtained and averaged.

Electrophysiological current traces were analysed with Minianalysis 6.0 (Synaptosoft Inc.) to extrapolate amplitude and onset time of all IPSCs or EPSCs recorded. The putative detection window for monosynaptic events was determined according to previously published criteria^41^. For each laser spot of the grid, all events whose onset fell within this detection window were summed and then averaged with values from different runs of the same experiment. Final heatmaps were built through a customized Matlab (Mathworks, US) script. In order to allow the reconstruction of the layer profile on the input map, a photomicrograph of the grids relative to the slice preparation was acquired and layer boundaries were manually determined. Normalized heatmaps were generated by dividing value in each spot by the sum of all pixels. Linear profiles (layer and columnar) were obtained by summing all values for each line in individual heatmaps. Average maps were obtained by aligning each individual map to the SP/Layer 6a boundary and averaging corresponding pixels.

### *In vitro* optogenetics stimulation

Optogenetics experiments were performed by conditionally expressing Channelrhodopsin 2 (ChR2, via the *Ai32* reporter allele) in SST+ interneurons (using *SST-ires-Cre*) interneurons or thalamic relay neurons (using *Olig3 Cre*). Wide-field light stimulation was delivered through a 40x objective to focus blue (470nm LED, CoolLED, UK) light onto the recorded cell. For each recorded SPN, two light stimulation duration pulses (1 and 10 ms) were employed at multiple LED power intensity to ensure that the minimal stimulation and full range of activation was captured irrespective of developmental age. For each LED pulse duration and intensity, five pulses were administered at a 20s interval and the evoked postsynaptic current (PSC) recorded.

Data analysis was performed through a customised Matlab script. Light-evoked PSCs were analysed if their onset was detected within 25 ms from the onset of the light stimulus; the relatively long latency was used to account for developmental effects that may affect ChR2 expression. For events within the mono-synaptic detection window, multiple PSC features were extracted such as amplitude, latency, 10-90% rise time, and decay time constant (τ). In particular, the latency of the PSC was calculated from the onset of LED stimulation; the decay τ was found by fitting a mono-exponential curve to the decay phase of the PSC; percentage of PSC occurrence was calculated throughout the five sweeps at each LED intensity.

In a subset of experiments, recordings of light-evoked IPSCs from SST+ interneurons were performed in a modified HDC ACSF containing 4 mM SrCl_2_ to replace CaCl_2_. Due to the slow onset of its effects, slices were bathed in Sr^2+^-containing HDC for a minimum of 20 min before light stimulation and data collection (Gil et al., 1999).

### Electrical stimulation of thalamic afferents

Thalamocortical (TC) afferent input to SPNs was tested using a bipolar microelectrode (Harvard Apparatus, UK) placed either in the ventrobasal nucleus (VB) of the thalamus or the internal capsule (IC) and connected to a current isolator (DS3, Digitimer Ltd, UK). The strength of the electrical stimulation was varied to find the minimal stimulation value^48, 49^, corresponding to EPSC evoked on ~50% of trials. The interstimulus interval was set at either 30 or 60s depending on developmental age. TC-EPSCs were considered if calculated standard deviation (jitter) of the EPSC latency was <1ms at minimal stimulation.

### Morphological reconstruction of recorded cells

Following electrophysiological assessment, slices containing biocytin-filled cells were fixed in 4% paraformaldehyde (PFA; diluted in phosphate-buffered saline, PBS) overnight at 4°C. Slices were then rinsed in PBS and incubated in 0.05% PBST containing Streptavidin-Alexa568 (1:500; Molecular Probes, US) for 48-72 h at 4°C. Slices were then washed 3x 10mins. in PBS and mounted on histology slides with Fluoromount (Sigma) mounting medium.

Slices were imaged through an Olympus FV1200 confocal microscope equipped with 10x or 20x dry objective. Z-stack images were acquired in order to maximize imaging of all neuronal processes containing biocytin to allow the offline morphological reconstruction. Image analysis was performed with Fiji-ImageJ software (NIH): confocal images of filled cells were selected for morphological reconstruction, performed using the Simple Neurite Tracer plugin. Dendrite directionality was calculated using the Directionality plugin implemented in Fiji-ImageJ onto reconstructed dendritic morphologies.

### Immunohistochemistry

Mice were terminally anaesthetized with pentobarbital (90 mg/kg) and transcardially perfused with 4% PFA in PBS. Dissected brains were incubated in PFA for 2 h at 4°C and then cryoprotected in 20% sucrose for 24 h at 4°C. Brains were then embedded into O.C.T. (VWR), frozen on dry ice and stored at −80°C. Each brain was sectioned into 14 to 16μm thick slices and mounted on histology slides; slides were stored at −20°C. Slides selected for immunohistochemistry were air-dried overnight at RT and washed 3x 10mins. at RT in PBS. Slides were then permeabilized for 30 min in 0.5% PBST (0.5 % Triton X-100 in PBS) and incubated in blocking solution (PBST 0.1%, Natural Goat Serum 5%) for 1 hour at RT. Primary antibodies used were chicken anti-GFP (ab13970, Abcam, dilution 1:250), rabbit anti-GABA (A2052, Sigma, 1:1000) and guinea-pig anti-GABA (ab17413, Abcam, 1:1500). Slides were incubated in primary antibody solution overnight at 4°C. Slides were then washed 3×10 min at RT in PBS and subsequently incubated in secondary antibody (Goat anti-Chicken IgG Alexa Fluor 488 conjugate, Goat anti-Rabbit IgG Alexa Fluor 568 conjugate; diluted in blocking solution 1:1000) for 2 hours at RT. Finally, the slides were washed in PBS, counterstained with DAPI (diluted 1:1000 in PBS) for 3 minutes at RT and mounted with Fluoromont (Sigma).

### Statistical analysis

All results are expressed as mean ± standard error of the mean; n indicates the number of cells recorded. Statistical analysis was performed with Prism (GraphPad, US). Normality and equal variance tests were run to direct the appropriate statistical test choice for comparison of parametric versus non-parametric datasets. Multiple groups were compared with a two-way ANOVA test; *post hoc* multiple comparisons were performed with the Holm-Sidak method. Two parametric groups were compared with Student’s t-test whereas two non-parametric groups were compared with the Mann-Whitney rank sum test. Finally, cumulative frequency distributions were compared with the two-sample Kolmogorov-Smirnov test. Statistical significance was evaluated at P ≤ 0.05; for the Kolmogorov-Smirnov test P ≤ 0.01 was considered statistically significant.

## Acknowledgements

Research in the Butt lab that contributed to this work was funded by the Medical Research Council (MRC)(MR/K004387/1), Biotechnology and Biological Sciences Research Council (BB/P003796/1), Human Frontiers Science Program Organisation (CDA0023/2008-C) and Brain and Behavior Research Foundation (Narsad; ref. 19079). Studentships awarded to FG and AM-S were funded by the Wellcome Trust; PGA was funded by an Imperial College London studentship; CV was funded by an MRC studentship. Funding for equipment came from the Wellcome Trust (089286/Z/09/Z) and OUP John Fell Fund (AV6721-C8000). Work in the Molnár laboratory related to early cortical circuit formation was funded by the MRC (G00900901, MR/N026039/1), Royal Society and Anatomical Society.

